# MYC Overexpression Confers Sensitivity to TACC3 Inhibition for Triple-Negative Breast Cancer

**DOI:** 10.64898/2026.04.28.721171

**Authors:** Mauricio Jacobo Jacobo, July Aung, Christopher O’Brien, Hope S. Rugo, Juha Klefström, Sourav Bandyopadhyay, Andrei Goga

## Abstract

The oncogenic transcription factor MYC is overexpressed across many cancer types, including triple-negative breast cancers (TNBCs), where it drives cell proliferation, chromosomal instability, and immune suppression. As MYC has proven difficult to target directly, identifying vulnerabilities created by MYC-driven mitotic rewiring is of significant therapeutic interest. Using a focused pharmacological screen of mitosis-related targets that are overexpressed in the context of MYC signaling, we identified several previously unrecognized dependencies on mitotic spindle proteins, including KIF18A and TACC3 in MYC^HIGH^ cells. We focused on TACC3 and found the small molecule inhibitor BO-264 induced pronounced micronuclei formation, cell cycle arrest, transcriptional reactivation of inflammatory signaling, and heightened cytotoxicity in MYC^HIGH^ models. TACC3 inhibition activates interferon-stimulated response element (ISRE) mediated inflammatory signaling and can activate the cGAS-STING pathway in a subset of triple-negative breast cancer contexts. Together, these findings identify TACC3 as a MYC-driven mitotic vulnerability. We highlight TACC3 inhibition as a promising strategy to impair tumor cell fitness and activate tumor-intrinsic inflammatory responses in MYC-driven cancers.

## Introduction

MYC is amplified or overexpressed in many of the most prevalent and difficult-to-treat human malignancies, including receptor triple-negative breast cancers (TNBC)^1^. As MYC regulates various biological processes, including cell-cycle progression, proliferation, apoptosis, and metabolic biosynthesis^2,3^, MYC profoundly shapes tumor behavior and therapeutic response. For instance, recent exploration of MYC-driven suppression of innate immune signaling has linked it to immune tolerance and reduced sensitivity to immune checkpoint blockade^4,5^. We and others have shown that MYC attenuates antitumor immunity by upregulating immune-evasive proteins such as CD47 (cluster of differentiation 47) and PD-L1 (programmed death-ligand 1)^6^, downregulating MHC class 1 (MHC-I) antigen presentation machinery^4^, and repressing interferon-related genes downstream of cGAS-STING signaling^5,7^. Despite its central role in tumor progression and immune evasion, MYC’s intrinsically disordered structure and lack of an established druggable pocket have resulted in a paucity of direct small molecule inhibitors^8^. These challenges underscore the need to identify MYC-dependent collateral vulnerabilities that can both limit tumor growth and bypass MYC-mediated immune suppression.

Mitotic inhibitors, which induce cytotoxicity by delaying mitosis and increasing chromosome segregation errors, offer a potential therapeutic approach for rapidly proliferating MYC^HIGH^ cells. Without therapeutic intervention, MYC overexpression leads to mitotic spindle defects, impaired chromosome alignment, and altered microtubule dynamics, resulting in delayed mitotic progression and increased chromosomal instability (CIN)^9,10^. Transcriptomic^9^ and proteomic^10^ analysis suggest that MYC-dependent rewiring of the mitotic proteome may help preserve mitotic progression despite the heightened spindle stress of MYC^HIGH^ cells. Upon pharmacological challenge with mitotic inhibitors, these cells exhibit amplified mitotic abnormalities and increased apoptosis, further underscoring MYC’s reliance on mitotic stress-buffering mechanisms^10,11^. This has motivated the development of MYC synthetic-lethal strategies targeting key mitotic regulators such as CDK1^1,12^ and Aurora B Kinase^13,14^. However, toxicity and off-target effects have limited their broader clinical utility^15–17^. Despite these limitations, low-dose strategies have been explored for their ability to elicit bystander killing by triggering inflammatory signaling downstream of severe chromosome segregation errors^10,18–25^. These observations suggest that selectively targeting mitotic proteins may provide a means to both exploit MYC-associated mitotic vulnerabilities and reactivate suppressed inflammatory signaling in MYC^HIGH^ tumors.

We hypothesized that perturbing mitotic spindle components in MYC^HIGH^ models would exacerbate chromosomal instability, impair cell division, and enhance cytotoxicity. To test this hypothesis, we conducted a focused small-molecule screen evaluating mitotic spindle inhibitors against genes whose expression is dysregulated in conditional MYC-expressing cell lines. This screen identified several synthetic-lethal dependencies, including a dependency on TACC3. Pharmacological inhibition of TACC3 produced clear on-target effects, including robust micronuclei formation and cell cycle arrest. Notably, TACC3 inhibition also triggered inflammatory signaling, although the specific pathways engaged varied across MYC^HIGH^ models. Together, our findings uncover a previously unrecognized MYC-dependent mitotic vulnerability with the potential to activate innate immune signaling and identify a subset of patients who may benefit from TACC3-directed therapies.

## Methods

### Cell culture and reagents

MTB/TOM cells were cultured using DMEM medium supplemented with 10% fetal bovine serum and 1% pen/strep as previously described^9^. Cells were maintained in a MYC^ON^ state with the addition of 1ug/ml doxycycline (Fisher #BP2653-5). To establish a MYC^OFF^ state, cells were withdrawn from doxycycline for 2 days prior to experimentation. Engineered human epithelial cell line RPE-NEO was previously described^12^ and grown in DMEM medium supplemented with 10% fetal bovine serum and 1% pen/strep. BT549 cells were obtained from the American Type Culture Collection (ATCC) and were grown according to published guidelines^91^. 4T1 cells were generously provided by Matthew Spitzer and maintained in RPMI-1640 medium supplemented with 10% FBS, 1% pen/strep, and 1% HEPES (CCFGL002, UCSF Cell Culture Facility). All cells were incubated at 37°C with 5% CO_2_. RPE-NEO cells were transduced with lentivirus containing the pCDH-puro-cMyc vector^92^ to overexpress MYC (RPE^MYC^). The pCDH-puro-cmyc vector was a gift from Jialiang Wang (Addgene plasmid # 46970; http://n2t.net/addgene:46970;). RPE-NEO cells transduced with an empty vector (CD510B-1, System Biosciences) were used as a control (RPE^CONTROL^). Cells were selected using 22 *u*g mL^-1^ puromycin for 11 days to establish stable RPE^MYC^ or RPE^CONTROL^ cell lines. To generate an ISRE reporter system, BT549 cells were transduced with lentivirus carrying the pGreenFire1-ISRE (EF1α-puro) (TR016PA-P, System Biosciences) construct and selected with 0.5 *u*g mL^-1^ puromycin for 10 days. All cell lines tested negative for mycoplasma contamination using PCR.

Paclitaxel (HY-B0015), Rabusertib (HY-14720), Kif15-IN-1 (HY-15948), MPI-0479605 (HY-12660), KHS101 Hydrochloride (HY-10996A), BTB-1 (HY-101770), AZ82 (HY-12241), and BO-264 (HY-135960) were purchased from MedChemExpress. diABZI (S8796), Vinorelbine (S4269), Monastrol (S8439), and Sovilnesib (E1180) were obtained from Selleck Chemicals. ABT-751 (A3169) was purchased from TCI America. Palbociclib (47284) was purchased from Cell Signaling. Eribulin was purchased from Eisai Europe (115457). All drugs were solubilized in DMSO and used at the indicated concentrations. Recombinant human IFNγ (11500-1) was obtained from PBL Assay Biosciences while recombinant mouse IFNγ (PMC4031) was purchased from Gibco.

### Immunofluorescence and micronuclei quantification

Cells were seeded on glass coverslips and fixed with 4% paraformaldehyde (PFA) in PBS for 10 min at room temperature. Cells were then permeabilized in 1X PBS/0.5% Triton X-100 for 15 min at room temperature. Nuclei were counterstained with DAPI (1;10,000) for 10min. Coverslips were then mounted using ProLong Gold Antifade mounting reagent (P36934, ThermoFisher) and imaged using an ECHO Revolution automated microscope equipped with a 20x/0.45 Fluorite LWD CC objective and accompanying imaging software. The images were further processed in ImageJ^93^. Micronuclei positive cells were calculated as a percentage of total cells per field. For quantification, 700-1000 cells per independent experiment were counted from multiple random fields. Micronuclei were defined as discrete DNA aggregates separate from the primary nucleus in cells where interphase primary nuclear morphology was normal. Mitotic and apoptotic cells were excluded from analysis.

### Transfection and siRNA treatment

Cells were reverse transfected using Lipofectamine RNAiMax Transfection Reagent (13778150, ThermoFisher), according to the manufacturer’s instructions. Non-targeting, mouse, and human TACC3 targeting siRNAs (D-001810-10-20, L-046198-00-0005, L-004155-00-0005) were purchased from Dharmacon (ON-TARGETplus SMARTPools, four siRNAs per gene). 50nM siRNA was used for all knockdown experiments.

### Cell viability assays

Cells were seeded in 96-well assay microplates overnight at various densities according to cell growth rate (for slow growing: cells ∼ 1000cells/well; for fast growing cells: ∼ 500cells/well), and then treated with drugs for 72 hrs. Cell proliferation and cell death were measured by staining with 1ug/ml Hoechst 33342 (Biotium, 40046) and 1ug/ml Propidium Iodide (ThermoFisher, P3566), respectively. Cells in stained plates were analyzed and nuclei counted using a BioTek Cytation 5 Cell Imaging Multimode Reader (Agilent) equipped with Gen5 imaging software. Cell proliferation was quantified from the total number of live cells, calculated by subtracting the number of PI-positive cells from the total number of cell nuclei. Cell death was assessed by determining the percentage of nuclei that were PI-positive. For drug screen analysis, AUC values were computed from baseline-subtracted PI measurements (ΔPI), where ΔPI = PI_Drug_ – PI_Vehicle_ for each DMSO-matched concentration. Drug sensitivity was compared between MYC^HIGH^ and MYC^LOW^ conditions using AUC fold-change (MYC^HIGH^ _AUC_/MYC^LOW^_AUC_), with ratios >1 indicative of heightened sensitivity in MYC^HIGH^ cells. For further validation of the effects of TACC3 inhibition or knockdown, cell viability was determined using the PrestoBlue HS Cell Viability Reagent (ThermoFisher, P50201) according to the manufacturer’s instructions. For colony outgrowth assays, cells were seeded in 12-well microplates at a density of 100-250 cells/well, allowed to adhere overnight, and then exposed to drug or DMSO for 9 d with medium change and drug refresh every 3-4 d. Cells were fixed with cold 100% methanol, stained with 0.5% crystal violet, and imaged using an HP Color LaserJet Pro MFP M479fdw scanner prior to quantification. For quantification, stains were extracted using an extraction buffer (0.1%SDS in 50% ethanol) and absorbance measured by a Tecan Safire2 plate reader at 590nm.

### Immunoblotting

Uncropped blots are provided (**Supplementary Fig. 5**). Cells for immunoblots were collected and lysed using Pierce RIPA buffer supplemented with protease (cOmplete Mini, EDTA-free, Roche) and phosphatase inhibitor cocktails (PhosSTOP, EASYpack, Roche) for 15 min on ice. Cell lysates were cleared by centrifugation at 14,000 r.p.m. for 10 min at 4°C. The supernatant was collected and protein quantified by BCA. Equal amounts of protein samples were resolved using Bolt 4-12% Bis-Tris Plus gels (Invitrogen) and transferred onto nitrocellulose membranes using the iBlot 2 Gel Transfer Device (Thermo). Quality control of the transfer was assessed by membrane staining with Ponceau S (Sigma Aldrich). Membranes were blocked and probed overnight on a 4°C shaker with primary antibodies (1:1,000 dilution unless indicated) recognizing the following proteins: TACC3 (ab134154, Abcam), TACC3 (8069, Cell Signaling), c-MYC (ab205818 HRP, Abcam, 1:10,000), p-Rb (Ser807/811) (8516, Cell Signaling), p21 (2947, Cell Signaling), Cyclin D1 (55506, Cell Signaling), Cyclin B1 (4138, Cell Signaling), p-Histone H3 (Ser10) (9701, Cell Signaling), p-TBK1 (Ser172) (5483, Cell Signaling), TBK1 (3504, Cell Signaling), p-IRF3 (Ser386) (37829, Cell Signaling), p-IRF3 (Ser396) (4947, Cell Signaling), IRF3 (11904, Cell Signaling), IRF3 (4302, Cell Signaling), STING (13647, Cell signaling), p-STAT1 (Tyr701) (9167, Cell Signaling), p-STAT2 (Tyr690) (PA5-97361, Thermo), STAT1 (14994, Cell Signaling), STAT2 (72604, Cell Signaling), Caspase 3 (9662, Cell Signaling), and β-actin (sc-47778 HRP, Santa Cruz Biotechnology, 1:10,000), GAPDH (5174, Cell Signaling, 1:10,000), Vinculin (13901, Cell Signaling, 1:10,000). Membranes were then incubated with horseradish peroxidase-conjugated secondary antibodies (1: 5,000 dilution) for 1 h at room temperature and developed using an enhanced chemiluminescence solution. Chemiluminescence was visualized using a ChemiDoc XRS+ imaging system equipped with Image Lab Software (Bio-Rad).

### Flow cytometric analysis

For cell cycle analyses, cells were seeded, treated as indicated, and harvested after 72h of incubation. Cells were fixed with cold 70% ethanol and stored at -20°C until the analysis day. On the same day of analysis, fixed cells were washed with PBS and resuspended with FxCycle PI/RNase staining solution (F10797, ThermoFisher). Samples were incubated for at least 30 minutes at room temperature and analyzed on a BD LSRII flow cytometer equipped with FACSDiva v9.0 software. Cell cycle calculations were performed using FlowJo v10.10.0.

For ISRE reporter activation assessment, BT549^ISRE-GFP^ reporter cell lines were treated with BO-264 or IFNγ for 72h. Cells were harvested, washed with PBS, and stained with LIVE/DEAD Fixable Near-IR dye (L10119, ThermoFisher, 1:1000) for 15 min at room temperature. Cells were washed with PBS and resuspended in FACS buffer for data collection using a BD LSRII instrument. The flow cytometry gating strategy, which involved gating on live cells and then on GFP positive cells (or ISRE-active cells) using untransduced BT549 as a negative control sample is shown in **Supplementary Fig. 3b**. Analysis of MHC cell surface expression was similarly performed except that harvested cells were stained with anti-human HLA-A/B/C conjugated to PE (560168, BD Pharmingen, 1:5) or anti-mouse MHC Class I conjugated to PE (12-5999-83, ThermoFisher, 1:100) for 25 min on ice prior to live/dead staining. Experiments were compensated with single color and unstained controls. Compensated cells were gated for FSC/SSC, singlets, and live cells. GFP positivity and MHC median fluorescence intensity were determined using FlowJo.

### Enzyme-linked immunosorbent assays

Cells treated with BO-264 for 72h were collected and processed as described previously. Intracellular 2’,3’-Cyclic GAMP (cGAMP) levels were quantified using the DetectX 2’,3’-cGAMP ELISA Kit (K067-H1, Arbor Assays) according to the manufacturer’s instructions. Normalized cGAMP was calculated by dividing the total amount of cGAMP detected in each sample by the corresponding total protein content. Alternatively, supernatants were collected from 72h treated BO-264 cells and cleared by centrifugation. CCL2 levels were quantified using either Human MCP-1/CCL2 ELISA Kit PicoKine (EK0441, Boster Bio) or Mouse MCP-1 ELISA Kit PicoKine (EK0568, Boster Bio) according to the manufacturer’s instructions. CXCL10 levels were quantified using Human CXCL10/IP-10 ELISA Kit PicoKine (EK0735, Boster Bio) or Mouse CXCL10/IP-10 ELISA Kit PicoKine (EK0736, Boster Bio) according to the manufacturer’s instructions. Normalized chemokine levels were calculated by dividing the total amount of chemokine detected in each sample by the corresponding total number of viable cells.

### Dendritic cell activation assay

Human buffy coats were obtained from Finnish Red Cross Blood Services under sample license number: 41/2025 for surplus blood products, from which PBMCs were isolated under provision for non-clinical research use as assessed by experts from the FRC Sample Service (https://www.veripalvelu.fi/en/sample-service/). CD14+ monocytes were isolated using positive magnetic separation according to the manufacturer’s protocol (Miltenyi Biotec) for in vitro generation of dendritic cells. For Mo-DC generation, isolated cells were cultured for 5 days in complete RPMI medium (31870074, Gibco) supplemented with 10% 56°C heat-inactivated (30’) FBS (25-079-CV, Corning), 100 U penicillin-streptomycin (15140-122, Gibco), 2mM L-Glutamine (25030-024, Gibco), 100ng/mL GM-CSF (130-093-864, Miltenyi Biotec) and 100ng/mL IL-4 (130-093-920, Miltenyi Biotec). Media was only refreshed once in between. On day 5, Mo-DCs were harvested and plated to 96-well U-bottomed plates (650180, Greiner-Bio) at densities of 30-100,000/100 uL depending on donor. Mo-DCs were subsequently activated for 24H by co-culture with 100 uL of conditioned media from BT549-treated cells. DCs were then stained (4°C, 30’) with CD11c (301604, BioLegend), HLA-DR (307606, BioLegend) and CD86 (305422, BioLegend) and washed thrice before analysis on Novocyte Quanteon and FlowJo v10.10.1. Cells with high expressions of HLA-DR and CD86 were considered activated.

### Analysis of public datasets

No new datasets were generated in this study. The MTB/TOM RNA-seq dataset was previously analyzed and published^9^ and downloaded from the Gene Expression Omnibus (GEO) repository (GSE130921). Batch-corrected, normalized gene expression data from cancer cell line models was obtained from the Cancer Dependency Map (DepMap) portal (https://depmap.org/portal), version 24Q4 (DepMap Public 24Q4)^29^. The mRNA z-score data, overall survival information, and ER/HER2/PR status of METABRIC^42^ and Breast Invasive Carcinoma TCGA PanCancer Atlas^94^ clinical datasets were obtained from cBioPortal for Cancer Genomics (https://www.cbioportal.org/)^27,28^. Normalized gene expression data and survival information from the I-SPY1^41^ trial were retrieved from the GEO repository under GSE22226. BO-264 (NSC 807620) drug sensitivity data (GI_50_) for the NCI-60 human cancer cell line panel were obtained from the NCI Developmental Therapeutics Program (DTP) drug screening database. Available at:https://dctd.cancer.gov/data-tools-biospecimens/data (accessed June 9, 2025). The GI_50_ values correspond to the concentration of compound required to inhibit cell growth by 50% as measured by the sulforhodamine B assay^38^. Gene set enrichment analysis (GSEA) of hallmark cancer gene signatures in the Molecular Signature Database version 2025.1.Hs^40^ was performed using GSEA version 4.4.0^95^. A MYC activity score was calculated by averaging the z-scored expression values of all genes in the HALLMARK MYC TARGETS V1 signature. Patients were stratified into low and high TACC3 expression groups using a cohort-specific threshold. In the METABRIC cohort, the lower and upper tertiles of TACC3 expression were used to define low and high groups, respectively, with the intermediate tertile excluded from analysis. In the I-SPY1 cohort, patients were dichotomized at the median TACC3 expression level. Survival was visualized using Kaplan-Meier survival curves, and associations between cohort groups and survival were assessed with Cox proportional hazards regression. All analyses were conducted in R using the “survival”and “survminer” packages.

### Statistical analysis

Data are expressed as means ± standard error of mean (±SEM), unless otherwise indicated. Statistical analyses were performed using GraphPad Prism 10 version 10.4.1 and R version 4.4.2. Two-tailed Student *t*-tests were used in all comparisons unless otherwise noted with *P* < 0.05 considered statistically significant throughout the study. All experiments were conducted independently at least 3 times.

### Data Availability Statement

All data generated or analyzed during this study are included in this published article and its supplementary information files. Cell lines generated in this study are available upon reasonable request from the authors.

## Results

### MYC Overexpression Enhances Sensitivity to TACC3 inhibition

We previously identified mitosis-related genes whose expression was altered in a MYC-dependent manner in a MYC-driven transgenic mouse model of triple-negative breast cancer (MTB/TOM), in which MYC expression is inducible with doxycycline^9,26^. We focused on genes significantly upregulated in MYC^ON^ cells compared to non-tumor tissue or when MYC is switched off (MYC^OFF^), also genes whose protein products are associated with regulation of mitotic processes, and for which small molecule inhibitors are readily available (**Fig. 1a**). Analysis of human breast cancer patient data from The Cancer Genome Atlas (TCGA)^27,28^ revealed that patients with a high MYC V1 gene-signature score, an assessment of MYC transcriptional activity, exhibited concurrent overexpression of these druggable mitotic genes (**Fig. 1b**). Similarly, assessment of mRNA expression profiles across multiple breast cancer cell lines from DepMap.org^29^ showed upregulation of these genes in cell lines with high MYC activity (**Fig. 1b**). Together, these findings indicate a strong association between elevated MYC activity and the expression of targetable mitotic regulators, many of which are critical for proper mitotic spindle formation and mitotic progression.

**Figure 1.**
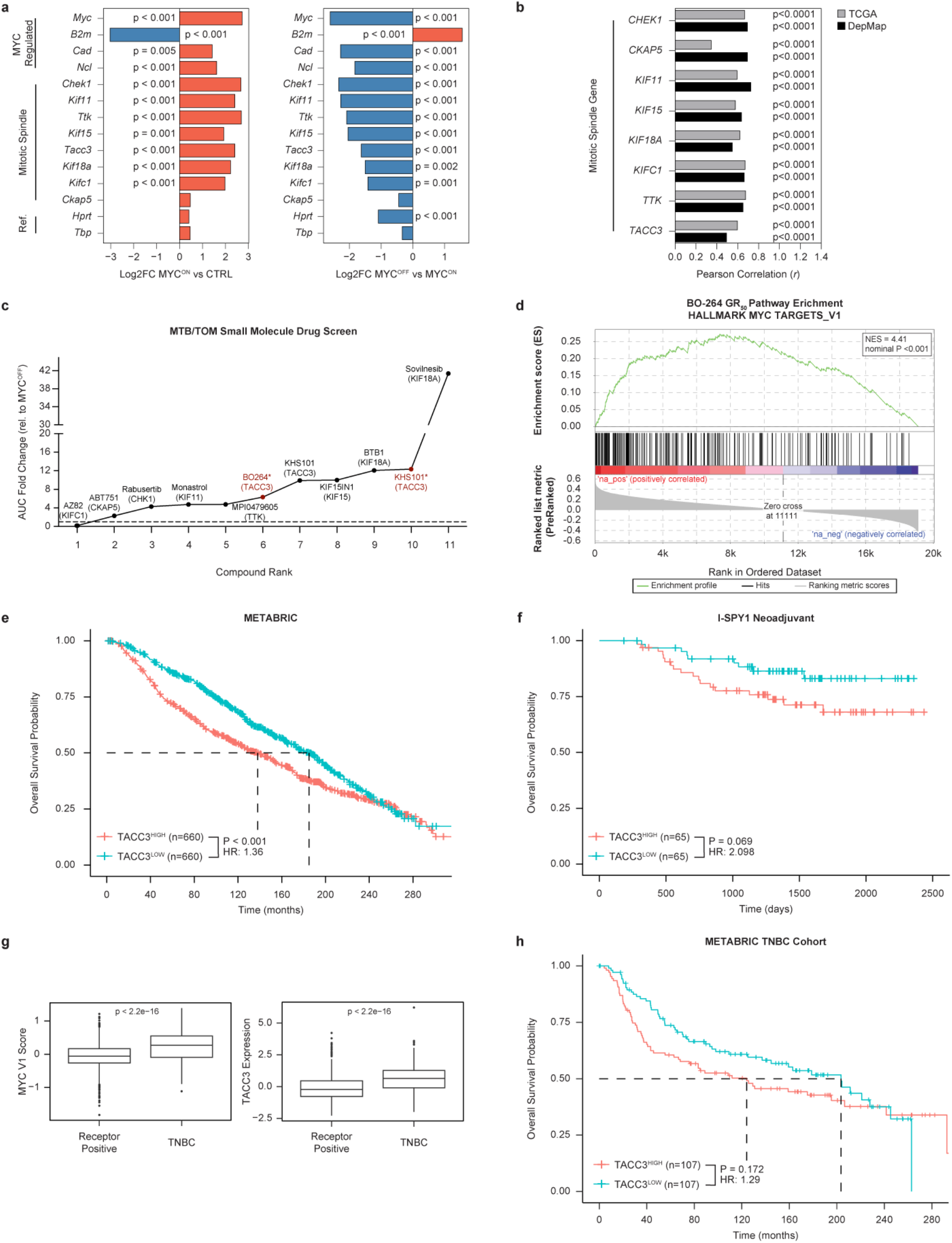
Heightened TACC3 expression in MYC high breast cancers is associated with worse survival. **a**, Differential expression analysis of mitotic spindle genes from previously published RNA-seq data in MTB/TOM tumors^9^. Left: MYC^ON^ tumors versus normal mammary tissue. Right: Tumors withdrawn from doxycycline for 3 days (MYC^OFF^) versus tumors grown in the presence of doxycycline (MYC^ON^). Adjusted *p*-value. **b**, Pearson correlation analysis of MYC V1 activity and deregulated mitotic spindle gene expression in the DepMap^29^ cell line (black) and TCGA breast cancer^27,28^ (gray) datasets. *P* value based on Pearson correlation. **c**, Plot showing compounds ranked by fold differences in cytotoxicity in MTB/TOM MYC^ON^ cells relative to MYC^OFF^ cells. **d**, Gene set enrichment analysis (GSEA) of a Hallmark MYC V1 Targets signature performed on a drug-response determined gene ranking list. NES, normalized enrichment score. **e**-**f**, Kaplan-Meier curves for overall survival in breast cancer patients. **e**, Neoadjuvant breast cancer I-SPY1 TRIAL a median TACC3 expression cutoff is used. **f**, METABRIC breast cancer trial TACC3 highest and lowest tertiles are plotted. **g**, Relative MYC V1 activity score and TACC3 expression in receptor-positive and triple-negative breast cancer tumors from the METABRIC cohort. *P* values are calculated using a two-sided Wilcoxon test. **h**, Kaplan-Meier curves for overall survival in TNBC patients from the METABRIC cohort. Breast cancer patients were dichotomized by TACC3 expression. Hazard ratios were estimated using Cox proportional hazards models.

Candidate targets included mitotic kinases, spindle proteins, kinesins, and microtubule regulators (**Table 1**). We screened MTB/TOM cells against a panel of anti-mitotic agents targeting the identified candidates (**Table 1**). MTB/TOM cells were cultured in the presence (MYC^ON^) or absence of doxycycline (MYC^OFF^) for two days before treatment with each inhibitor at four concentrations (80nM - 40μM) for 72 hrs. The relative cytotoxicity in MYC^ON^ versus MYC^OFF^ cells was calculated for each compound. In this screen, anti-mitotic agents targeting CHK1 (Rabusertib), Eg5 (Monastrol), KIF18A (BTB-1; Sovilnesib), and TACC3 (KHS101) exhibited several-fold greater cytotoxicity in MYC^ON^ cells (**Fig. 1c**). Enhanced sensitivity to CHK1 and Eg5 inhibition is consistent with known MYC-associated vulnerabilities^10,11,30,31^, supporting the validity of the screening approach. Notably, TACC3 and KIF18A emerged as less characterized MYC-dependent targets. While KIF18A inhibition has been proposed as a therapeutic strategy for cancers with chromosomal instability^32,33^, TACC3 inhibition has been linked to tumors with centrosome amplification^34^. Given less is known about the function of TACC3 in MYC^HIGH^ cancers, we sought to further evaluate TACC3 dependency. We next tested sensitivity to BO-264, a more potent and selective TACC3 inhibitor^35^, in a secondary screen. Notably, BO-264 is more selective for TACC3 than KHS101^36,37^. BO-264 treatment, like KHS101, demonstrated increased sensitivity of MYC^ON^ cells to TACC3 inhibition (**Fig. 1c**). Next, we sought to identify gene expression patterns associated with BO-264 sensitivity. We correlated individual gene expression levels with BO-264 GI_50_ values across multiple cancer types using the NCI-60 human cancer cell line panel.^38^ The resulting correlations were ranked to generate a drug-response-associated gene signature.^39^ Gene set enrichment analysis (GSEA) of this ranked signature revealed significant enrichment of the Hallmark MYC V1 signature^40^ (**Fig. 1d**). These data indicate that BO-264 sensitivity correlates with MYC transcriptional activity, with MYC^HIGH^ cells showing greater sensitivity to TACC3 inhibition across multiple cancer cell types. Taken together, these findings suggest TACC3 as a novel vulnerability in MYC^HIGH^ cancers.

**Table 1.**
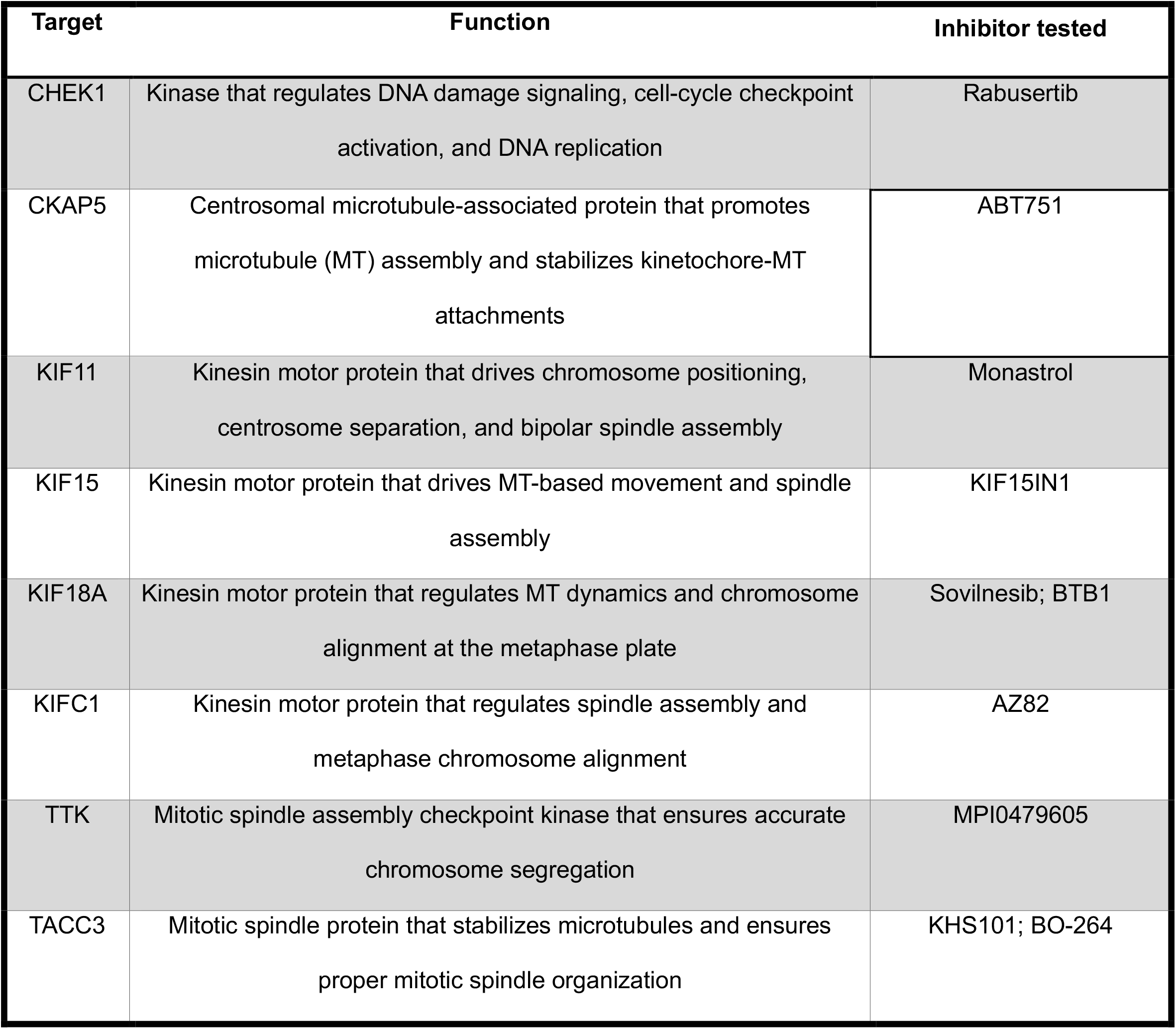
Candidate Mitotic Spindle Targets.

To assess the clinical relevance of TACC3 in MYC^HIGH^ cancers, we analyzed breast cancer patient outcomes in the I-SPY1^41^ and METABRIC cohorts.^42^ Across all breast cancer cases, high TACC3 expression (TACC3^HIGH^) was associated with worse overall survival (I-SPY1: HR = 2.09, p = 0.069; METABRIC: HR = 1.36, p < 0.001) (**Fig. 1e-f**) and increased likelihood of relapse (I-SPY1: HR = 1.48, p = 0.217; METABRIC: HR = 1.58, p < 0.001) (**Supplementary Fig. 1a-b**). We subsequently examined TNBC cases within METABRIC, a subset that frequently exhibits MYC overexpression and we found also has elevated TACC3 expression (**Fig. 1g**). Among TNBC patient subtype, TACC3^HIGH^ tumors showed a trend toward poorer overall survival and recurrence-free survival compared to TACC3^LOW^ tumors (OS: HR = 1.29, p = 0.172; RFS: HR = 1.47, p = 0.064) (**Fig. 1h, Supplementary Fig. 1c**). Although some associations did not reach statistical significance, the consistent magnitude and direction of effect observed across both cohorts and survival endpoints suggest an adverse role for elevated TACC3 expression in MYC^HIGH^ breast cancers, supporting its further investigation as a potential therapeutic vulnerability in this aggressive subtype.

Having demonstrated TACC3’s potential as a MYC-associated therapeutic target, we next sought to obtain a comprehensive drug response profile for BO-264, assessing its anti-proliferative and cytotoxic effects in MYC^LOW^ and MYC^HIGH^ contexts. To this end, we tested the short-term effects of BO-264 in two isogenic cell line models: MTB/TOM cells with inducible MYC expression^4,9^ and RPE1 cells which we engineered to stably overexpress MYC. Notably, induction of MYC expression in both systems led to elevated TACC3 protein levels (**Supplementary Fig. 2a**). In MTB/TOM cells, BO-264 treatment suppressed proliferation and increased the abundance of the apoptotic marker cleaved caspase-3 in MYC^ON^ cells compared to MYC^OFF^ cells (**Fig. 2a-b**). Similarly, RPE cells overexpressing MYC exhibited decreased proliferation (**Fig. 2c**) and viability (**Fig. 2d**) with in response to BO-264 treatment compared to control RPE cells. To further evaluate a genetic dependence on TACC3 in MYC high cells, we depleted TACC3 with a pool of siRNAs in both MTB/TOM and RPE cells, confirming knockdown efficiency by western blot (**Supplementary Fig. 2b**). TACC3 depletion significantly increased cell death in both MYC^LOW^ and MYC^HIGH^ contexts, but had a more pronounced effect in MYC^HIGH^ cells (**Fig. 2e**). As MYC overexpression is associated with increased cellular proliferation, we performed growth rate (GR) inhibition analysis to normalize drug sensitivity for differences in proliferation rate between MYC^LOW^ and MYC^HIGH^ cells^43,44^. GR_50_ analysis confirmed that the increased sensitivity of MYC^HIGH^ cells to BO-264 is independent of growth rate differences (**Supplementary Fig. 2c**). Next, we sought to determine the long-term effects on cellular proliferation using a colony formation assay. RPE^MYC^ cells demonstrated reduced long-term clonogenic survival compared to RPE^CONTROL^ control cells (S**upplementary Fig. 2d-e**). Altogether, these results demonstrate that TACC3 inhibition preferentially compromises the survival of MYC-overexpressing cells.

**Figure 2.**
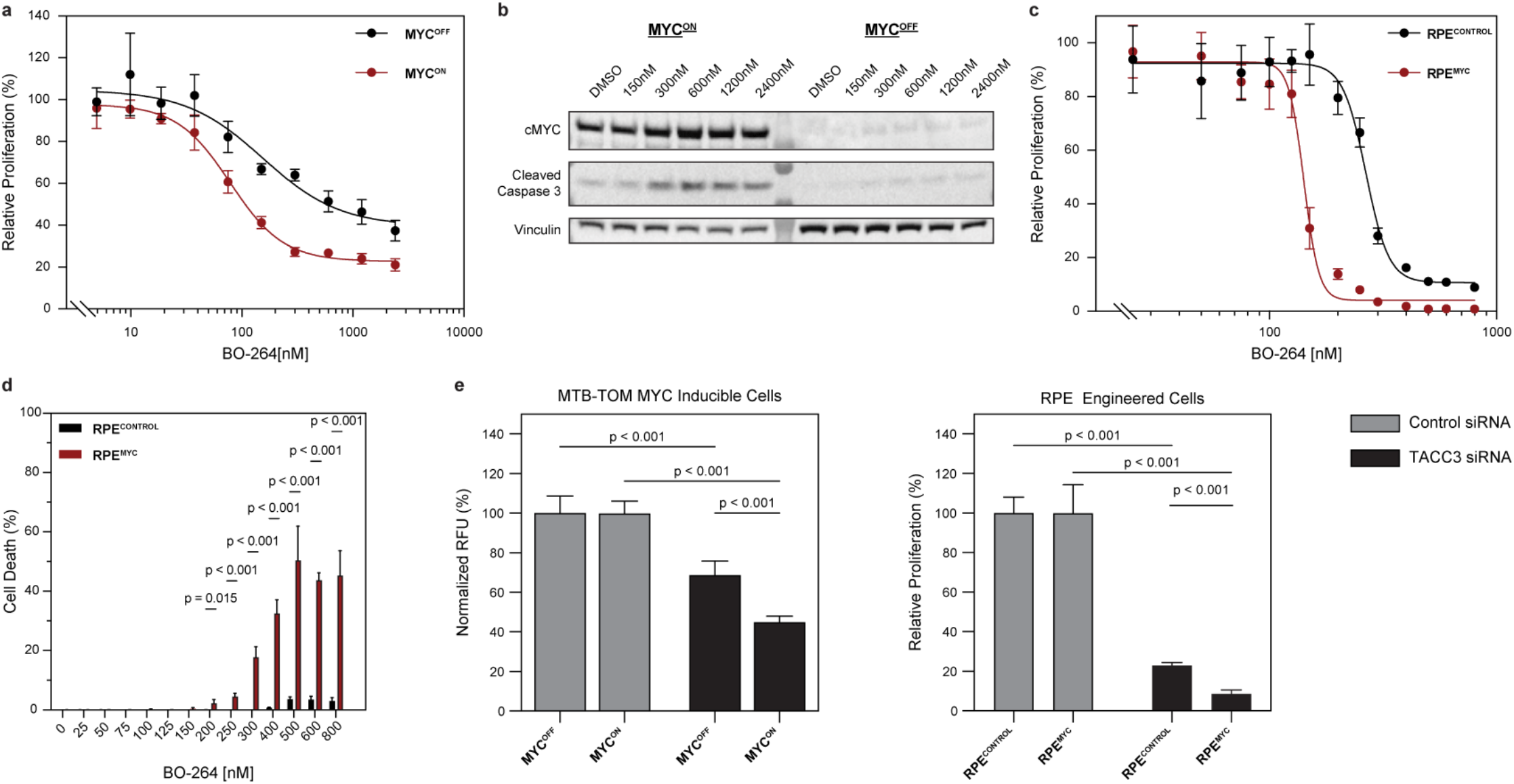
MYC overexpressing cancers display increased sensitivity to TACC3 inhibition. **a**, Proliferation of MYC^ON^ and MYC^OFF^ MTB/TOM cells in response to 72 h BO-264 treatment, normalized to their respective vehicle controls. **b**, Immunoblot analysis of MYC^ON^ and MYC^OFF^ MTB/TOM cells treated with BO-264 for 72 h to test the effect on an apoptosis marker. Vinculin is shown as a loading control. Representative image from *n* = 3 independent experiments. **c**, Proliferation of RPE^CONTROL^ and RPE^MYC^ cells in response to 72 h BO-264 treatment, normalized to their respective vehicle controls. **d**, Fraction of total cells undergoing cell death by PI staining in RPE^CONTROL^ or RPE^MYC^ cells treated with BO-264 for 72 h at indicated concentrations. **e**, Proliferation of MYC^OFF^ or MYC^ON^ MTB/TOM cells (left) and RPE^CONTROL^ or RPE^MYC^ cells (right) in response to 72 h TACC3 siRNA knockdown, normalized to their respective non-targeting siRNA controls. Error bars are mean ± s.d., and *p* values calculated using a two-sided *t*-test; * P < 0.05, *** P < 0.001, **** P < 0.0001. For (**a**) and (**c)**, data show a representative experiment with *n* = 4 technical replicates (*n*=10 for **e**). Each experiment was repeated independently three times, with similar results observed.

Small-molecule inhibitors, such as KHS101, have been proposed to act by reducing TACC3 stability^45^. Therefore, we sought to determine whether BO-264 similarly promotes TACC3 destabilization. Following BO-264 treatment, we observed reduced TACC3 protein abundance in RPE^MYC^ and BT549 TNBC cells compared to DMSO controls (**Supplementary Fig. 2f**). Co-treatment with the proteasome inhibitor MG132 prevented loss of TACC3 protein expression (**Supplementary Fig. 2f**), suggesting that BO-264 promotes proteasome-mediated degradation of TACC3. These data indicate that BO-264, like other published TACC3 inhibitors^45^, induces protein destabilization, providing a mechanistic basis for the growth inhibition and cytotoxic effects observed above.

### TACC3 inhibition induces micronuclei formation in MYC high cells

Genetic loss of TACC3 causes defective kinetochore-microtubule attachments and abnormal mitotic spindle morphology, leading to chromosome misalignment and micronuclei formation^34,46,47^. Formation of micronuclei is a hallmark of chromosomal instability limiting mitotic fidelity. We tested if BO-264 treatment or siRNA-mediated knockdown of TACC3 for 72 hours could elicit increased micronuclei formation. BO-264 treatment led to a significant increase in cells with micronuclei in multiple cell lines in which MYC is overexpressed, including engineered (RPE^MYC^ cells), human TNBC (BT549 cells), and a mouse TNBC model in which MYC is highly expressed (4T1 cells), but not in MYC^LOW^ RPE^CONTROL^ cells (**Fig. 3a-b**). Similarly, depletion of TACC3 with a pool of siRNAs induced micronuclei accumulation across all cell models, with more pronounced and statistically significant effects in TNBC models BT549 and 4T1 cells (**Fig. 3c**). Collectively, these data demonstrate that BO-264 phenocopies TACC3 genetic depletion in inducing genomic instability.

**Figure 3.**
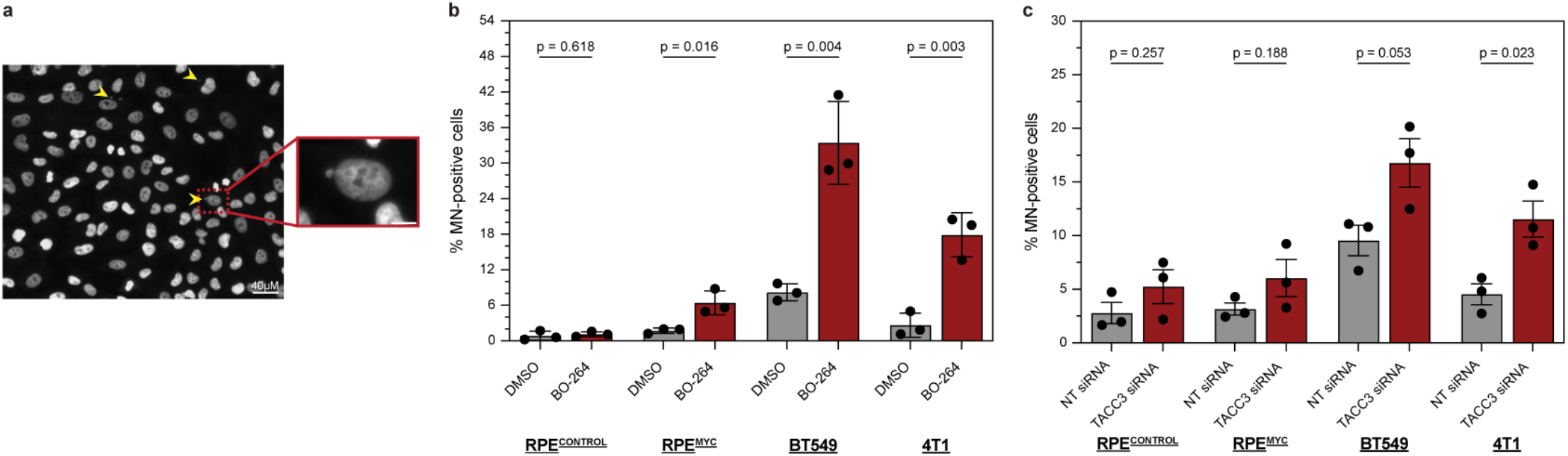
TACC3 inhibition induces mitotic defects. **a**, Representative image of RPE^MYC^ cells treated with 167nM BO-264 for 72 h with DAPI staining DNA. Arrows indicate micronuclei. Higher magnification of RPE MYC cells positive for micronuclei shown below. Scale bars 10uM unless indicated otherwise. **b-c**, Percentage of micronucleated RPE^CONTROL^, RPE^MYC^, BT549, and 4T1 cells 72 h after treatment with vehicle or BO-264 (**b**) or transfection with non-targeting (NT) or TACC3 siRNA (**c**). Data is an average of multiple high-powered (20x) fields analyzed per sample (700-1000 total cells/sample). Data representative of *n* = 3 independent experiments. Error bars are mean ± s.e.m., and *p* value calculated using a two-sided t-test.

### Cell Cycle Disruption is More Pronounced in MYC high cells

Given that TACC3 perturbation induces micronuclei formation, and mitotic errors are frequently associated with cell cycle arrest in the subsequent G1-phase^48–50^, we investigated whether TACC3 inhibition altered cell-cycle distribution. TACC3 has been implicated in regulating both mitotic progression and G1/S transition in cancer cells^34,35,46,51^, potentially reflecting its diverse interactome, which includes proteins involved in centrosome function as well as a role for overcoming G1/S checkpoint control^34^. TACC3 knockdown in non-tumorigenic RPE cells led to increased G1 accumulation and reduced cyclin D1 and phosphorylated RB expression, consistent with G1 arrest (**Fig. 4a-b**). Likewise, BO-264 treatment in RPE^MYC^ but not RPE^CONTROL^ cells increased the G1 population, upregulated the CDK inhibitor p21, and decreased phosphorylated RB, consistent with G1 arrest (**Fig. 4a-b, Supplementary Fig. 3a**). The minimal cell cycle effect of BO-264 in control RPE^CONTROL^ cells is consistent with their reduced sensitivity to drug exposure.

**Figure 4.**
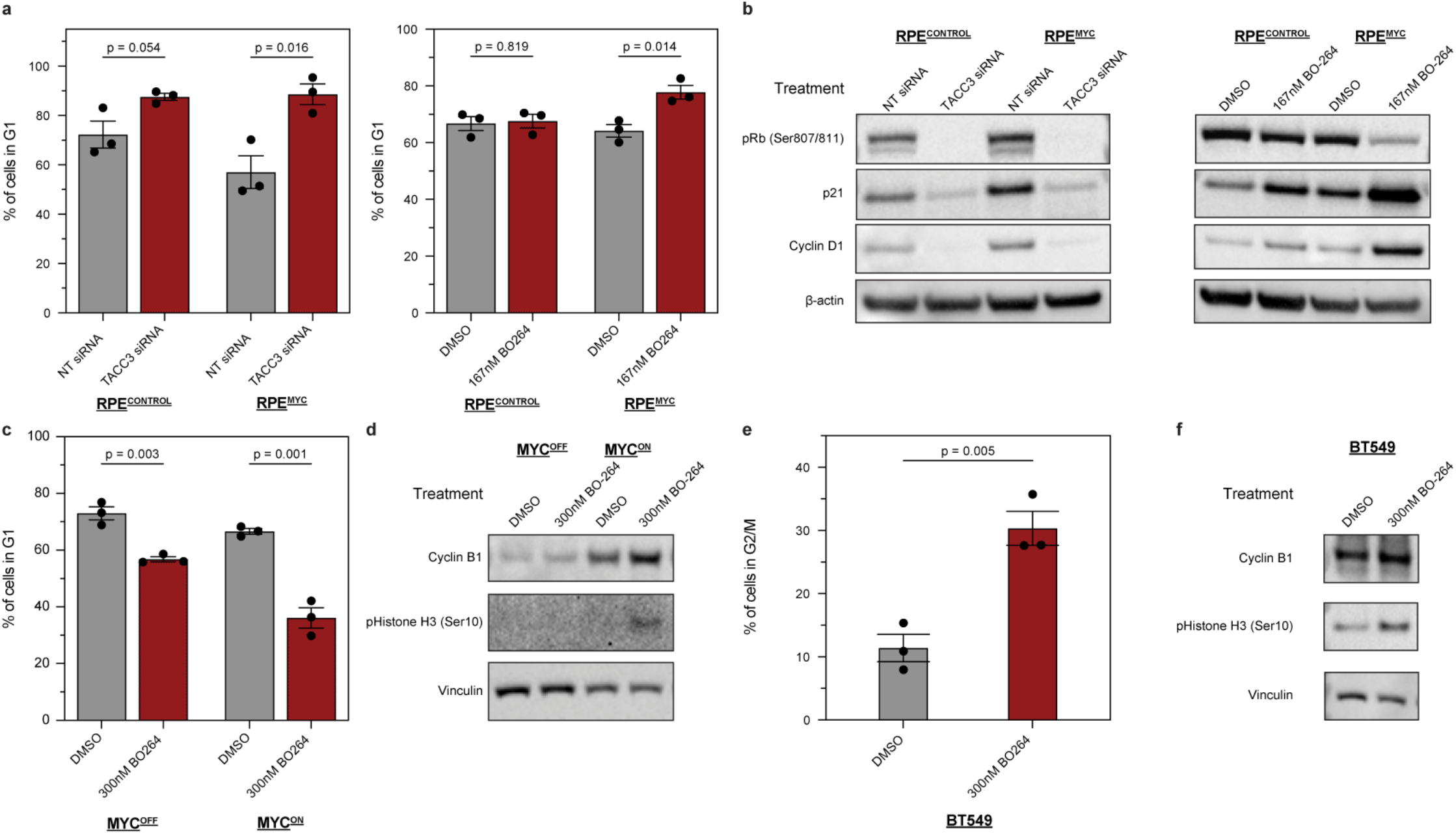
TACC3 inhibition disrupts cell cycle progression. **a**, Flow cytometry cell cycle analysis with PI staining in RPE^CONTROL^ and RPE^MYC^ cells transfected with TACC3 siRNA (left) or treated with 167nM BO-264 (right) for 72 h. **b**, Immunoblot analysis of G1/S progression markers in RPE^CONTROL^ and RPE^MYC^ cells transfected with TACC3 siRNA (left) or treated with 167nM BO-264 (right) for 72 h. β-actin is shown as a loading control. Representative image from *n* = 3 independent experiments. **c**, Flow cytometry cell cycle analysis with PI staining in MYC^OFF^ and MYC^ON^ MTB/TOM cells treated with 300nM BO-264 for 72 h. **d**, Immunoblot analysis of mitotic arrest markers in MYC^OFF^ and MYC^ON^ MTB/TOM cells treated with 300nM BO-264 for 72 h. Representative image from *n* = 3 independent experiments. **e**, Flow cytometry cell cycle analysis with PI staining in BT549 cells treated with 300nM BO-264 for 72 h. **f**, Immunoblot analysis of mitotic arrest markers in BT549 cells treated with 300nM BO-264 for 72 h. Representative image from *n* = 3 independent experiments. Data representative of *n* = 3 independent experiments. Error bars are mean ± s.e.m., and *p* values calculated using a two-sided *t*-test.

Unlike non-tumorigenic engineered RPE1 cells, ∼80-90% of TNBCs harbor TP53 mutations or loss^52,53^, which are broadly associated with altered G1/S checkpoint control^54,55^. To assess whether this genomic context might influence cell-cycle responses to BO-264, we examined cell-cycle arrest across MYC^HIGH^ TNBC models: human BT549 TNBC cells and MYC^ON^ MTB/TOM cells. In contrast to the G1 arrest observed in RPE cells, MYC^ON^ MTB/TOM cells exhibited mitotic arrest with 4N DNA content. BO-264 treatment decreased the fraction of cells in G1 phase (**Fig. 4c**) while increasing cyclin B1 expression and phospho-Histone H3 (Ser10) (**Fig. 4d**), markers of M arrest, in MYC^ON^ compared to MYC^OFF^ cells. As with RPE^CONTROL^ cells, BO-264 had minimal effects in MTB/TOM MYC^OFF^ cells, further supporting MYC-dependent TACC3 vulnerability. BO-264 treatment of human BT549 TNBC cells resulted in mitotic arrest with increased 4N DNA accumulation and prominent induction of phospho-Histone H3 (Ser10) (**Fig. 4e-f, Supplementary Fig. 3a**). Notably, BT549 cells exhibited markedly higher micronuclei formation compared to RPE^MYC^ cells, correlating with the more pronounced G2/M accumulation. Collectively, these data confirm TACC3 as a key regulator of cell cycle progression, with the arrest phase varying depending on cellular contexts.

### TACC3 inhibition upregulates inflammatory signaling

MYC overexpression elicits down-regulation of MHC-I and induction of other immune suppressive pathways^4,5,7^. Nevertheless, inflammatory responses can be triggered through interferon-independent induction of interferon-stimulated genes (ISGs), driven by the detection of mislocalized nucleic acids by pattern recognition receptors (PRRs)^56,57^. Notably, micronuclei contain unincorporated chromosome fragments encapsulated by a fragile nuclear envelope. Upon rupture, these DNA fragments can enter the cytoplasm, where they stimulate DNA-damage pathways and can be recognized by PRRs such as cyclic GMP-AMP synthase (cGAS), leading to downstream ISG transcription^18^. Having established that BO-264 treatment significantly increases micronuclei formation (**Fig. 3b)**, we sought to determine whether this chromosomal instability could elicit an inflammatory response. To test whether BO-264-induced genotoxic stress drives inflammatory signaling, we employed multiple complementary approaches, including measurement of ISG reporter activity, upregulation of MHC class I surface expression, chemokine secretion, and activation of nucleic acid-sensing PRR pathways.

We first examined ISG expression that is initiated by transcriptional activation of interferon-stimulated response element (ISRE)-containing genes^58,59^. To monitor ISRE activity, we engineered TNBC BT549 cells to express a destabilized GFP reporter under ISRE control, hereafter referred to as BT549^ISRE-GFP^ cells (**Fig. 5a**). Untransduced BT549 cells were used to define the threshold for GFP positivity (**Fig. 5b, Supplementary Fig. 3b**). BT549^ISRE-GFP^ cells exhibited low baseline GFP expression under control conditions (**Fig 5b-c**). As proof of principle, treatment of cells with IFNγ, which robustly activates interferon signaling, led to strong GFP expression (**Fig. 5b-c, Supplementary Fig. 3b**). Notably, TACC3 inhibition with BO-264 for 72 hrs also significantly induced GFP expression (**Fig. 5b-c**). These results demonstrate that BO-264 activates ISRE-driven transcription in BT549 cells, indicating engagement of inflammatory signaling pathways.

**Figure 5.**
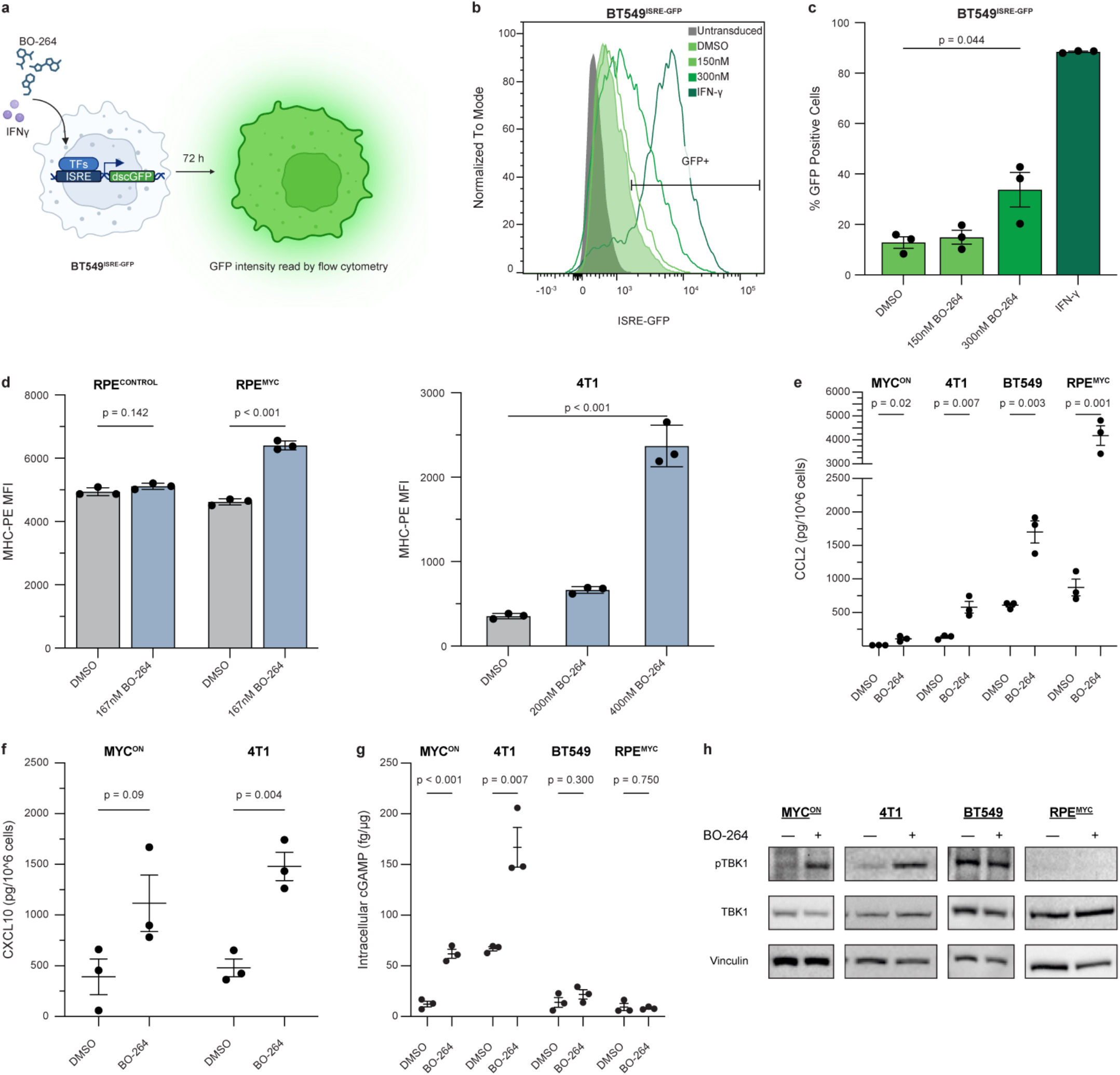
BO-264 displays immunostimulatory abilities in MYC-overexpressing cells. **a**, Diagram of IFN induction assay: BT549 cells contain an IFN reporter expressing destabilized GFP (dscGFP). ISRE: IFN-stimulated response elements. GFP median fluorescence intensity (MFI) was assessed 72 h after treatment by flow cytometry. Created with BioRender.com. **b**, Representative flow cytometry plot depicting GFP MFI of BT549^ISRE-GFP^ cells treated with vehicle, BO-264, or IFNγ for 72 h. Untransduced BT549 cells were used to establish GFP-positivity. **c**, Quantification of GFP positivity in BT549^ISRE-GFP^ cells after 72 h treatment. **d**, Flow cytometry analysis of MHC MFI in RPE (left) and 4T1 (right) cells treated with BO-264 for 72 h. Representative experiment with three samples per condition shown. Error bars are mean ± s.d., and *p* values calculated using a two-sided *t*-test. Trends repeated across three independent cell passages. (**e**-**f**), ELISA analysis of secreted CCL2 (**e**) and CXCL10 (**f**) levels in the indicated cell lines following treatment with vehicle or BO-264 for 72 h. **g**, ELISA analysis of intracellular cGAMP levels in the indicated cell lines upon treatment with vehicle or BO-264 for 72 h. **h**, Immunoblot analysis of cGAS-STING pathway activity in MTB/TOM MYC^ON^, 4T1, BT549, and RPE^MYC^ cells treated with BO-264 for 72 h. β-actin is shown as a loading control. Representative image from *n* = 3 independent experiments. Data representative of *n* = 3 independent experiments. Error bars are mean ± s.e.m., and *P* values calculated using a two-sided *t*-test.

Our second approach was to evaluate MHC class 1 (MHC-I) gene expression and cell surface expression. MHC-I and processing-associated genes are typically induced by interferon signaling^60^ but are suppressed by high MYC expression^4^ (**Fig. 1a**). In the process of mounting an immune response, these MHC-I molecules help facilitate antigen presentation and cytotoxic T-cell recruitment^61–63^. To determine whether TACC3 inhibition restores MYC-driven suppression of antigen presentation, we measured MHC-I surface expression in BO-264-treated RPE and 4T1 cells. Following 72 h treatment with BO-264, we observed a significant increase in MHC-I protein on the cell surfaces of RPE^MYC^ and 4T1 cells (**Fig. 5d**). Interestingly, RPE^CONTROL^ cells showed no change in MHC-I surface expression, consistent with their reduced sensitivity to BO-264 (**Fig. 2c, 5d**). These data indicate that MYC’s repression of antigen presentation machinery can be bypassed by TACC3 pharmacological inhibition.

We took a third approach to evaluate inflammatory signaling induced by TACC3 inhibition; we extended analysis beyond tumor-intrinsic outputs to non-cell-autonomous effects that drive the secretion of chemokines with immunomodulatory potential^64,65^. We tested whether BO-264 treatment in MYC^HIGH^ cells induces secretion of immune cell modulators. We quantified levels of the myeloid cell attractant CCL2^66^ and the effector T-cell chemoattractant CXCL10^67^ in cell supernatants. BO-264 treatment induced robust CCL2 secretion across all four MYC^HIGH^ cell models, consistent with induction of inflammatory signaling (**Fig. 5e**). While CXCL10 secretion was predominantly observed in BO-264-treated 4T1 cells, with a modest increase in MTB-TOM MYC^ON^ cells (**Fig. 5f**). Both BT549 and RPE^MYC^ cells showed no detectable CXCL10 secretion, suggesting that release of inflammatory cytokines is dependent on additional cellular characteristics. Release of CCL2 and CXCL10 suggest an inflammatory chemokine profile favoring myeloid cell recruitment over effector T-cell attraction.

A fourth approach assessed whether BO-264-induced inflammatory signaling translated into functional immune modulation, including processes associated with T-cell priming^68^. We performed immunophenotyping of dendritic cells exposed to conditioned-medium from treated tumor cells. CD14^+^ monocytes isolated from peripheral blood mononuclear cells (PBMCs) were differentiated into immature monocyte-derived DCs and subsequently co-cultured with conditioned medium from BT549 cells treated with either vehicle, BO-264, eribulin, vinorelbine, or paclitaxel. Eribulin and vinorelbine, are mitotic inhibitors previously found to promote dendritic cell maturation^22^, and were included as positive controls. Paclitaxel, a mitotic inhibitor shown to exhibit weak dendritic cell maturation^22^, was included as a negative control. Consistent with prior reports, eribulin and vinorelbine induced dendritic cell activation in samples from five out of six donor PBMCs, whereas paclitaxel and BO-264 failed to increase activation relative to vehicle controls across all donors tested (**Supplementary Fig. 4**). Thus, while BO-264 elicits tumor-intrinsic inflammatory signaling and chemokine secretion, these responses may be insufficient to license productive antigen-presenting cell activation in the BT549 TNBC cells. Notably, BT549 highly expresses MYC, which inhibits CCL5 secretion as well as STAT1 expression and phosphorylation in these cells^7^. Thus, while all models tested demonstrated CCL2 secretion, CXCL10 release and MHC-I upregulation were cell model-dependent.

Lastly, we investigated the mechanism underlying inflammatory signaling in BO-264-treated cells. We hypothesized that inflammatory signaling could be mediated by cGAS-STING pathway activation downstream of genomic instability^18^. In support of this model, we observed elevated levels of the STING activator cyclic guanosine monophosphate-adenosine monophosphate (cGAMP) in BO-264-treated mouse tumor models MTB/TOM MYC^ON^ and 4T1 cells (**Fig. 5g**). In addition, BO-264-treated MTB/TOM MYC^ON^ and 4T1 cells exhibited increased levels of phosphorylated TBK1, consistent with activation of the cGAS-STING signaling axis (**Fig. 5h**). Notably, BO-264-treated human cells RPE^MYC^ and BT549 cells showed no increase in cGAMP levels or TBK1 phosphorylation (**Fig. 5g-h**). These results indicate that cGAS-STING signaling contributes to inflammatory responses in a subset of BO-264-treated MYC^HIGH^ cell models. However, the absence of cGAS-STING activation in RPE^MYC^ and BT549 cells, despite robust ISRE activation in BT549 (**Fig. 5b-c**) and MHC-I upregulation in RPE^MYC^ (**Fig. 5d**), suggests that alternative PRR pathways may mediate inflammatory signaling in these contexts. This lack of cGAS-STING engagement in BT549 and RPE^MYC^ cells following BO-264 treatment may partially account for the absence of CXCL10 secretion and DC maturation. Collectively, these findings demonstrate the presence of heterogeneous inflammatory signaling programs induced by BO-264 treatment across MYC^HIGH^ models, resulting in distinct immune-stimulatory phenotypes engaged by both STING-dependent and STING-independent inflammatory pathways.

## Discussion

MYC activation is a hallmark of tumor initiation and maintenance, driving extensive transcriptional reprogramming that supports uncontrolled proliferation, genomic instability, and immune evasion^69–72^. Its deregulation profoundly alters mitotic fidelity, creating dependencies on core mitotic machinery that can be therapeutically exploited^9–11^. Given MYC’s undruggable nature, indirect strategies that target MYC-induced vulnerabilities have gained considerable attention. Our findings build on this concept by identifying previously unrecognized dependencies on mitotic spindle-associated genes in MYC^HIGH^ contexts through pharmacological inhibition. In addition to confirming established MYC-dependent vulnerabilities such as Eg5 and CHK1, we uncovered novel sensitivities to TACC3 and KIF18A inhibition, and to a lesser extent other mitosis-associated druggable targets (**Fig. 1c; Table 1**). We prioritized TACC3 for in-depth characterization given that multiple inhibitors we tested had strong cytotoxic effects at nanomolar concentrations. TACC3 inhibition exacerbated mitotic dysfunction, manifested by micronuclei formation and cell cycle arrest in MYC^HIGH^ cells, ultimately leading to heightened cytotoxicity. The screen identified KIF18A as another MYC-associated vulnerability and further validation and mechanistic characterization of KIF18A inhibition may reveal additional therapeutic opportunities and further define the landscape of targetable mitotic dependencies in MYC-driven cancers. Collectively, these results reinforce the therapeutic potential of targeting MYC-driven mitotic dependencies and establish TACC3 inhibition as a promising therapeutic strategy.

TACC3 is a mitotic spindle protein that regulates microtubule stability, microtubule dynamics, and centrosome integrity^34,46,73– 76^. Comprehensive analysis has shown that TACC3 is distinctly upregulated across multiple tumor types compared to normal tissues, and its elevated expression correlates with poor survival in breast and other cancers (**Fig 1e-f, 1h, Supplementary Fig. 1a-c**) ^35,77–79^. Here, we demonstrated that TACC3 is overexpressed at the transcript level in MYC^HIGH^ tumors and cancer cell line models. MYC overexpression in non-tumor epithelial cells or MYC conditional transgenic models of TNBC increased TACC3 protein levels, suggesting that TACC3 expression is positively regulated by MYC activity. Consistent with these findings, breast cancer patients with elevated TACC3 expression exhibited worse overall and recurrence-free survival. Although it remains unclear whether MYC directly regulates TACC3 transcription, the correlation between MYC and TACC3 expression suggests potential regulatory mechanisms. Previous studies have identified FOXM1, a known MYC target gene^80^, as an upstream activator of TACC3^34^. One plausible mechanism is that TACC3 expression could be indirectly regulated by MYC through FOXM1, though this remains to be experimentally validated in these models. Regardless of the precise regulatory mechanisms, TACC3’s association with high MYC activity and poor clinical outcomes supports its use as an exploratory biomarker for MYC-driven cancers.

TACC3 contains a highly conserved C-terminal domain that mediates multiple protein-protein interactions critical for centrosome organization and microtubule stabilization during mitotic progression^34,73–76^. Under MYC^HIGH^ conditions associated with rapid proliferation, cells experience heightened mitotic stress and instability, creating a dependency on mitotic spindle components to ensure faithful chromosome segregation^9–11^. The observed increase in micronuclei formation and cell cycle arrest upon TACC3 inhibition likely reflects an amplification of mitotic destabilization in MYC-overexpressing cells. Mechanistic studies have shown that BO-264 directly interacts with the C-terminal TACC domain, establishing BO-264 as one of the most potent and selective TACC3 inhibitors available^35^. Here, we demonstrate that BO-264 also induces proteasome-mediated degradation of TACC3 in MYC^HIGH^ cell models. In future studies, immunoprecipitation-based analyses of TACC3 interaction networks disrupted by BO-264 in MYC^HIGH^ contexts would be valuable to explore. Collectively, these findings indicate that TACC3 supports mitotic fidelity in the context of MYC overexpression, and that its perturbation exposes the fragile balance of mitotic control in MYC-driven cancers. Further investigation into TACC3’s protective role through overexpression studies in MYC^HIGH^ cells could clarify its function in maintaining mitotic progression and chromosomal stability in this context.

The heightened chromosomal instability observed following TACC3 inhibition in MYC^HIGH^ cells suggested potential engagement of innate immune signaling pathways. Under MYC^HIGH^ conditions, inflammatory responses are typically suppressed, resulting in a poorly immunogenic cell state that limits immune cell infiltration, tumor recognition, and tumor cell clearance^4,6,7^. This suppression has been shown to contribute to the limited efficacy of immune checkpoint blockade in MYC-driven breast cancers^4,5,81,82^. Cytosolic DNA derived from ruptured micronuclei can activate the cGAS-STING pathway, eliciting tumor cell-intrinsic inflammatory signaling^18,83^. Consistent with this mechanism, we observed elevated cGAMP production and downstream TBK1 phosphorylation following TACC3 inhibition in mouse TNBC models MTB/TOM MYC^ON^ and 4T1 cells, which reinforces the link between mitotic disruption and immune pathway activation. However, activation of cGAS-STING signaling was not uniform across models. For example, BT549 cells, which displayed robust ISRE activation and micronuclei formation, showed no detectable cGAS-STING signaling. Similarly, TACC3 inhibition in RPE^MYC^ cells stimulated increased MHC-I cell surface expression despite lacking STING pathway activation. These findings highlight the complexity of inflammatory signaling following TACC3 inhibition, likely reflecting cell-type-specific engagement of distinct but overlapping pattern recognition receptor pathways that sense aberrant nucleic acid species and act cooperatively or dominantly to induce interferon-stimulated gene (ISG) expression^57,84^. This may reflect differential activation of DNA-sensing pathways, such as cGAS-STING or TLR9, as well as RNA-sensing pathways mediated by RIG-I, MDA5, or TLR3, depending on the nature of the aberrant nucleic acids generated and the repertoire of PRRs expressed. Future studies should determine whether this heterogeneity in innate immune signaling arises from differences in PRR engagement^85^ or from epigenetic^86^ and transcriptional regulation of nucleic acid-sensing pathways and downstream interferon signaling programs^7,87^.

Chemokine profiling and functional immune cell phenotyping revealed that TNBC models had distinct responses. For example, there was a consistent induction of CCL2 secretion across all of the cell models suggesting that TACC3 inhibition could lead to myeloid cell recruitment. In contrast, CXCL10 secretion varied across models, and similar to paclitaxel treatment, induction of dendritic cell maturation was not observed. While myeloid populations are frequently associated with immunosuppressive functions in tumors^88^, they may also contribute to tumor cell clearance under certain conditions^89^. Future studies should define how BO-264-induced cytokine and chemokine release reshapes the tumor immune microenvironment and immune cell functionality, and whether these changes promote anti-tumor immunity or immune suppression. Additional functional assays, such as cell-based phagocytosis assays, may provide further insight into the mechanisms of clearance of tumor cells in response to BO-264. Collectively, these findings demonstrate that TACC3 inhibition bypasses MYC-mediated immune suppression and induces inflammatory signaling in MYC^HIGH^ tumors. However, this tumor-intrinsic inflammation may not be sufficient in all breast cancers to stimulate effective anti-tumor T-cell responses and may require additional combination immune strategies.

Among the limitations of this study, we did not perform *in vivo* evaluation of BO-264 tumor response in MYC-driven breast cancers. Although BO-264 is currently the most potent TACC3 inhibitor available, its poor pharmacokinetic properties have limited its translational potential^90^. Nonetheless, previous studies have shown that BO-264 significantly reduced tumor growth in MDA-MB-231 xenografts and in the TNBC patient-derived xenograft (PDX) model TM01278, without observable toxicity^34^. Given that TNBC exhibits high MYC expression in the majority of TNBC patients^1^, including in MDA-MB-231 cells^1^, the observed antitumor effects in these models support the therapeutic potential of TACC3 inhibition in MYC-driven contexts. Furthermore, because previously published xenograft studies were conducted in immunocompromised models, it remains critical to determine whether the cell-intrinsic inflammatory responses observed *in vitro* translate into meaningful anti-tumor immunity using immunocompetent models. Notably, A2A Pharmaceuticals has developed AO-252, a next-generation TACC3 inhibitor with improved pharmacological properties that is currently undergoing clinical evaluation for advanced solid tumors (clinicaltrials.gov; NCT06136884) but was not available for our preclinical studies. Future preclinical and clinical studies using such optimized compounds will be essential to assess the therapeutic potential of targeting TACC3 in MYC-driven breast cancers. Nevertheless, our work supports stratification of breast cancer patients with high MYC activity as candidates who may benefit from TACC3-targeted therapies.

## Author Contributions

M.J.J., A.G. and H.S.R. contributed towards study conceptualization. M.J.J. designed and performed experiments and data analysis supporting this study. C.M.O. conducted colony outgrowth assays. J.A. performed dendritic cell-based functional validation and analysis. M.J.J and A.G. wrote the manuscript with input from all authors. A.G., S.B., and J.K. supervised the studies.

## Acknowledgements

We thank members of the Goga and Klefström laboratories, Sourav Bandyopadhyay, and Alejandro Sweet-Cordero for helpful discussions and technical assistance. This work was supported by NCI U01CA168370 (S.B.), NIGMS R01GM107671 (S.B.), CDMRP DoD W81XWH-21-1-0774 (A.G.), CDMRP DoD W81XWH-21-1-0773 (J.K.), NIH R01CA266756 (A.G.), Subramanian Breast Cancer Support Fund (A.G., H.S.R), Breast Cancer Research Foundation (H.S.R., A.G.), Mark Foundation Endeavor Award (A.G.), Gazarian Family Foundation (A.G.), and Prospect Creek Foundation (S.B., A.G.). We acknowledge the PFCC (RRID: SCR_018206) and the Biomedicum Flow Cytometry Core Facility (RRID: SCR_024470) for assistance in generating flow cytometry data. Support was also received from the Finnish Red Cross Blood Service (license number: 41/2025). The schematic of the interferon induction assay in Figure 5A was generated using BioRender (Goga, A. [2026] https://BioRender.com/94v6×99).

## Supplemental Information

### Supplemental Figures and Legends

**Supplementary Figure 1.**
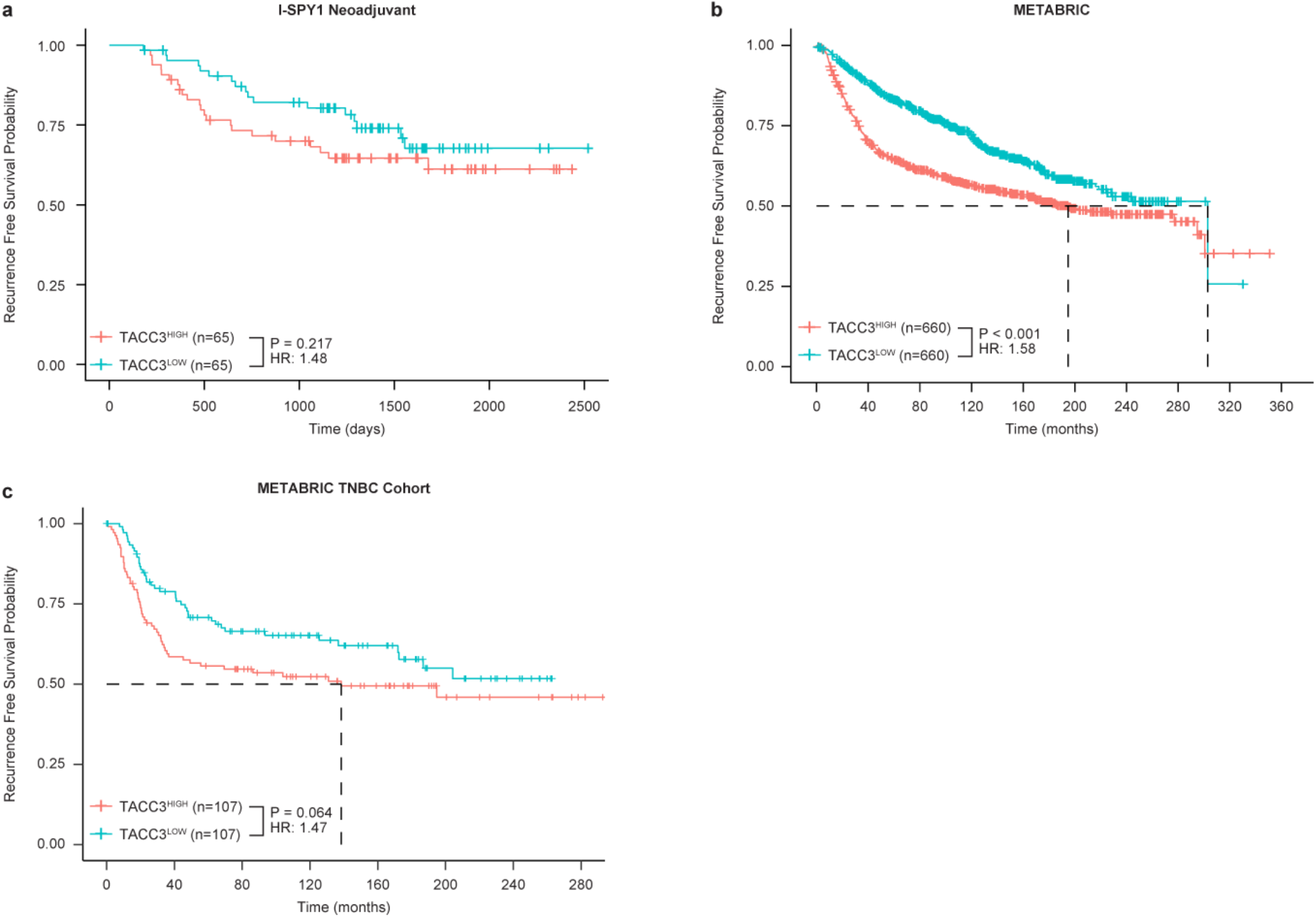
Recurrence-free survival (RFS) in TACC3^HIGH^ breast cancer patients. Kaplan-Meier curves for RFS were generated after dichotomizing I-SPY1 (**a**) and METABRIC (**b**) cohorts by TACC3 expression. METABRIC breast cancer patients were further stratified by receptor status, and RFS was evaluated within the TNBC subtype (**c**). Hazard ratios were estimated using Cox proportional hazards models.

**Supplementary Figure 2.**
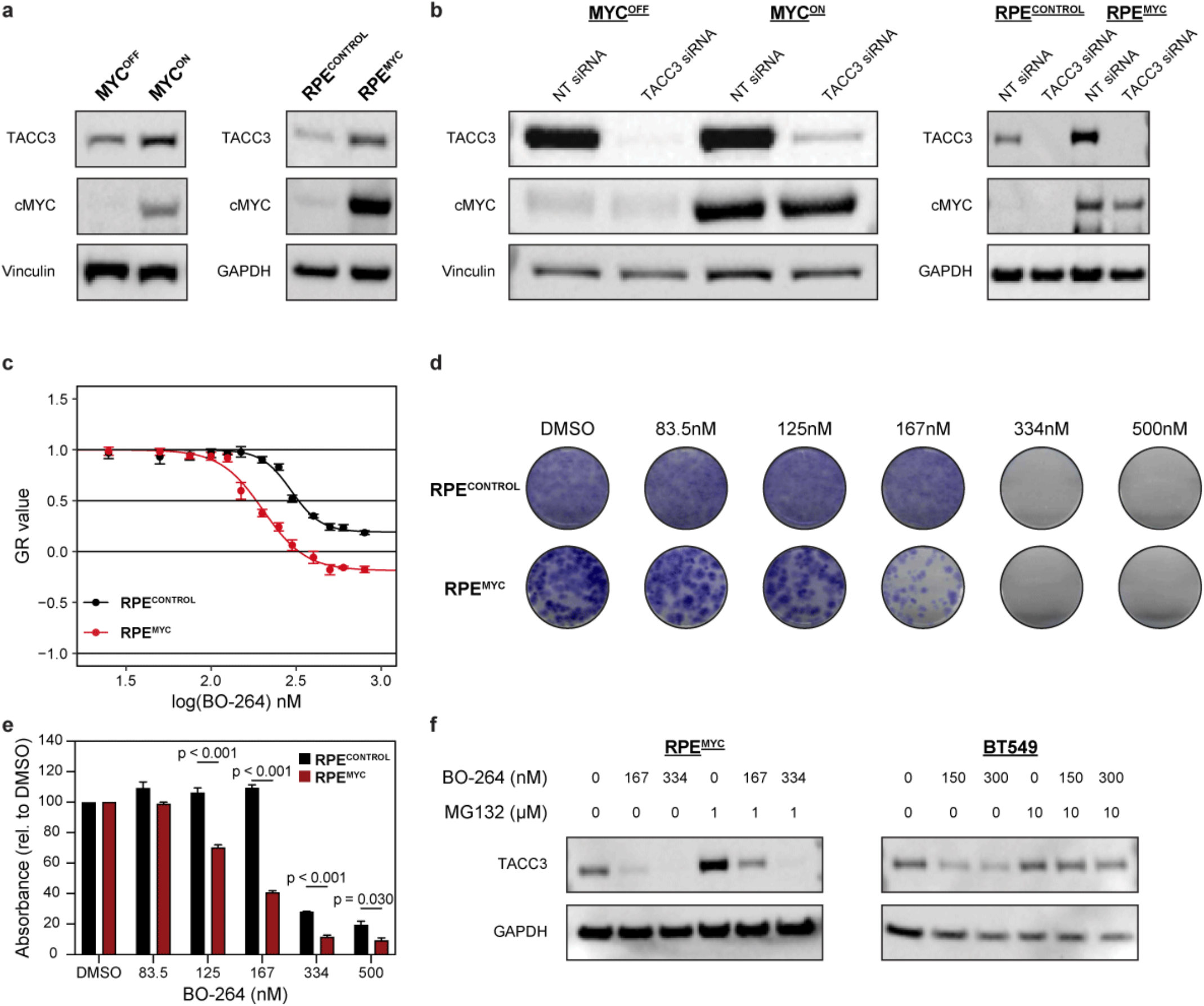
BO-264’s antiproliferative effects in MYC^HIGH^ cells. **a**, Immunoblot analysis of TACC3 expression in MTB/TOM and RPE cells. Representative image from *n* = 3 independent experiments. **b**, Immunoblot validation of siRNA knockdown of TACC3 in MYC^OFF^ and MYC^ON^ MTB/TOM cells (left) and RPE^CONTROL^ and RPE^MYC^ cells (right). Representative image from *n* = 3 independent experiments. **c**, Growth rate inhibition measurements of RPE^CONTROL^ and RPE^MYC^ cells in response to 72 h BO-264 treatment. Data show a representative experiment with *n* = 4 technical replicates. Error bars are mean ± s.d., and *p* values calculated using a two-sided *t*-test. Trends repeated across three independent cell passages. **d**, Crystal violet staining of RPE^CONTROL^ and RPE^MYC^ cells after 9 d treatment with increasing concentrations of BO-264. Images are representative of *n* = 3 independent experiments with similar results. **e**, Quantification of crystal violet stained RPE^CONTROL^ and RPE^MYC^ cells in response to various doses of BO-264. Error bars are mean ± s.e.m. *P* value was calculated using a two-sided *t*-test; * P < 0.05, *** P < 0.001, **** P < 0.0001. **f**, Immunoblot analysis of TACC3 proteasome-mediated degradation in RPE^MYC^ (left) and BT549 (right) cells treated with BO-264 for 72 h. Prior to harvesting, cells were additionally incubated in the presence or absence of MG132. GAPDH is shown as a loading control. Representative image from *n* = 3 independent experiments.

**Supplementary Figure 3.**
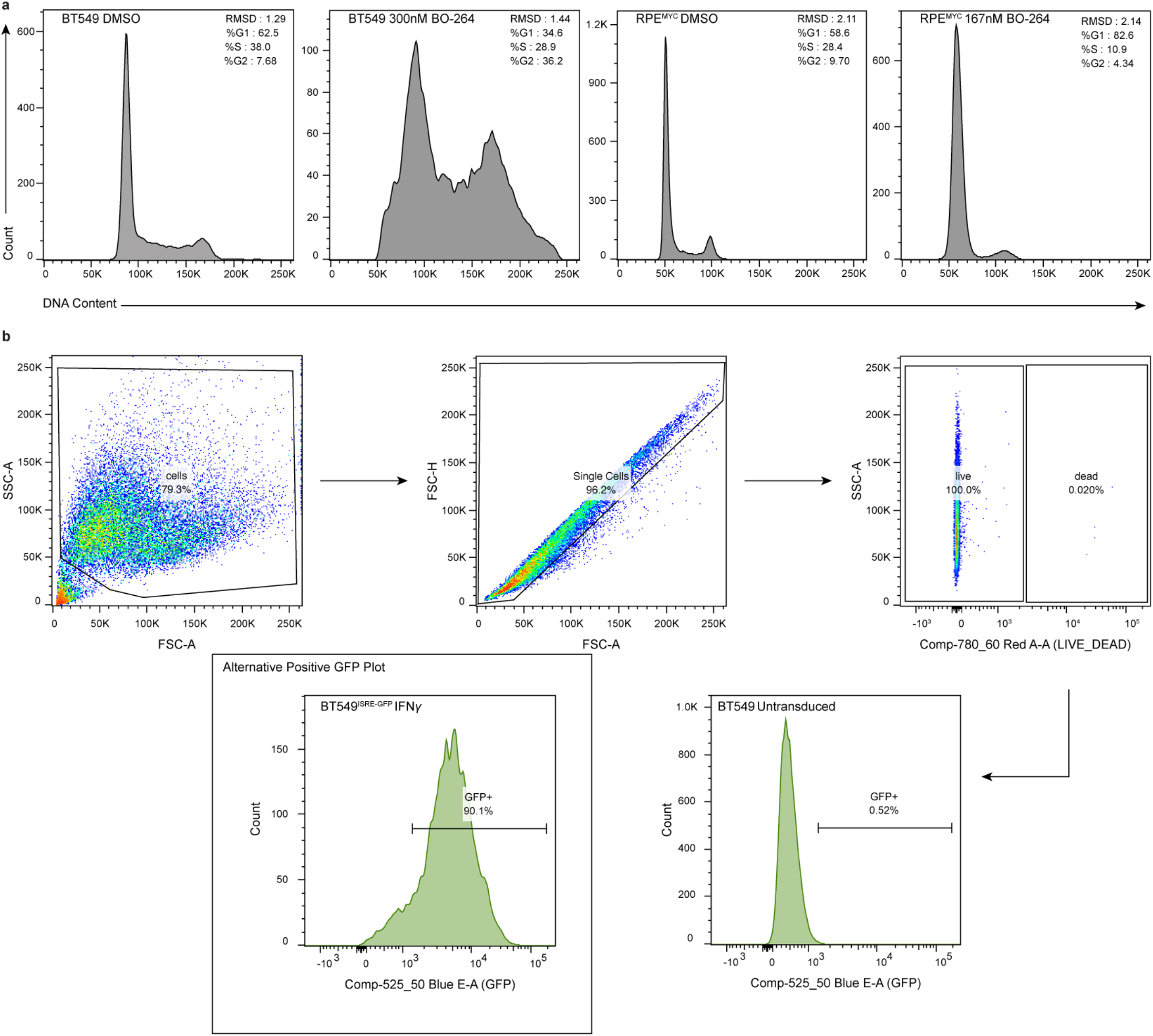
Flow cytometry gating strategy to analyze live cell populations. **a**, Representative cell cycle profiles of RPE^MYC^ and BT549 cells after treatment with BO-264 for 72 h. The percentage of cells in G1, S, and G2-M phases of the cell cycle, as determined by DNA content based on propidium iodide staining, is indicated. **b**, Flow cytometry gating strategy to identify GFP-positive cells (ISRE-active cells). Representative experimental control sample, untransduced BT549 cells. Side scatter and forward scatter were used to exclude debris. Forward scatter was used to distinguish single cells. Live cells were identified as negative for live/dead staining. GFP-positive live cells were identified by setting the fluorescence threshold based on untransduced BT549 controls. For reference, GFP signal intensity from live BT549^ISRE-GFP^ cells treated with IFNγ is shown to the left.

**Supplementary Figure 4.**
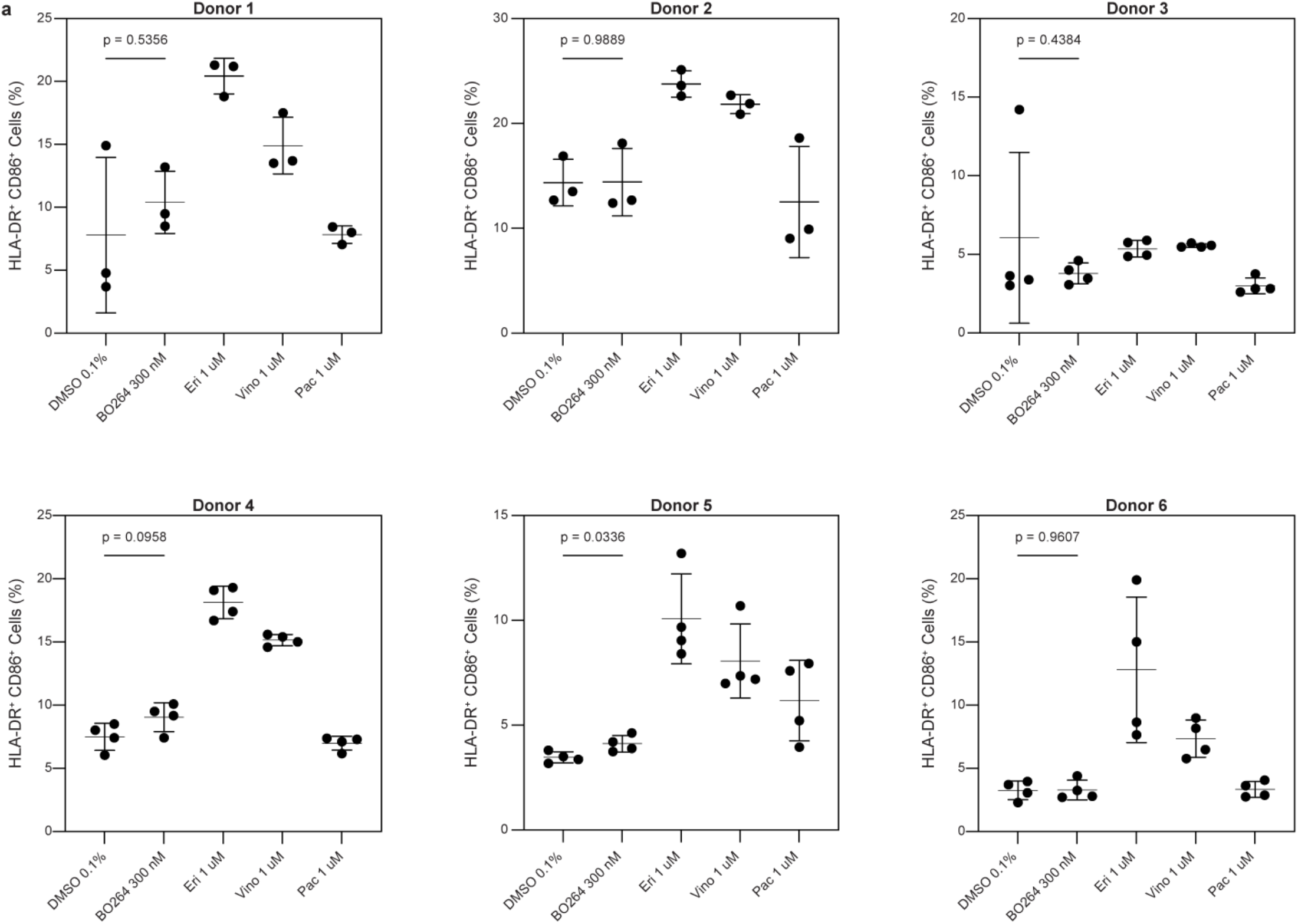
Dendritic cell (DC) functional assays reveal poor DC maturation following BO-264 treatment. **a**, Flow cytometry analysis of DC activation (HLA-DR^+^ CD86^+^) across six donors following DC co-culture with conditioned medium from BT549 cells treated with indicated drugs for 72h. Data are representative of *n* = 3 technical replicates per donor. Error bars indicate mean ± s.d., and *P* values were calculated using a two-sided *t*-test.

**Supplementary Figure 5.**
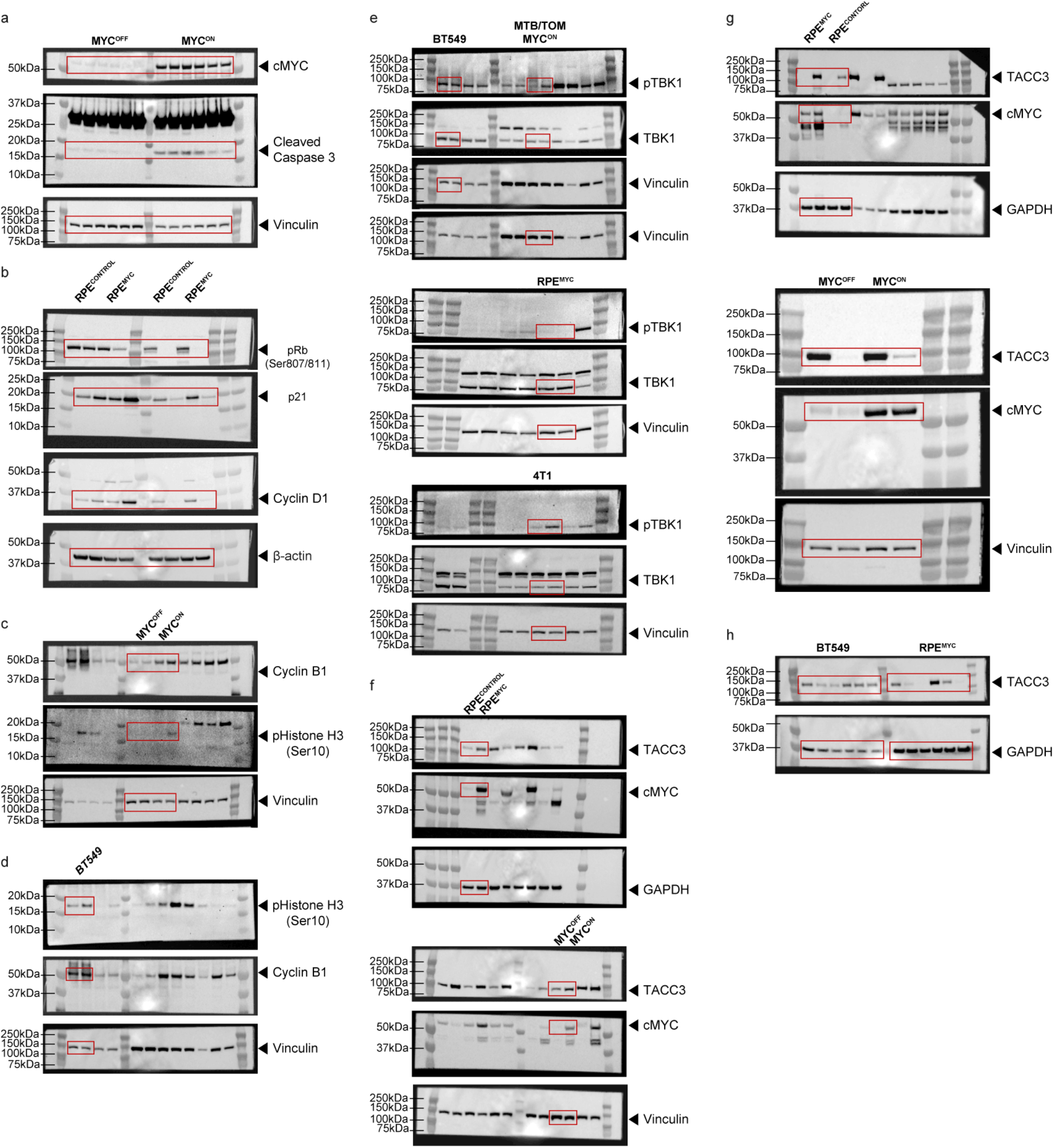
Full blot images. Regions surrounded by red boxes are presented in (**a**) Fig. 2b, (**b**) Fig. 4b, (**c**) Fig. 4d, (**d**) Fig. 4f, (**e**) Fig. 5h, (**f**) Supplementary Fig. 2a, (**g**) Supplementary Fig. 2b, and (**h**) Supplementary Fig. 2f.

